# Cnidarian oocytes reveal conserved actin-driven mechanisms of female meiosis

**DOI:** 10.64898/2026.09.16.751942

**Authors:** Yamini Vadapalli, Komal Bhattacharyya, Sascha Lambert, Bishal Samanta, Antonio Z. Politi, Stefan Klumpp, Peter Lenart

## Abstract

Oocyte meiosis is a specialized form of cell division adapted to produce a single haploid egg for fertilization. Diverse actin-driven mechanisms have essential roles in supporting these highly asymmetric divisions of the large oocyte. Whether these represent exotic adaptations in individual species or whether actin has broadly conserved functions in animal oocytes remains unclear. To address this, we established live-imaging assays, combined with targeted perturbations and biophysical modeling, in the non-bilaterian jellyfish *Clytia hemisphaerica*. We show that in *Clytia,* a nuclear F-actin network stabilizes the large oocyte nucleus. Upon meiotic entry, a transient F-actin shell forms to facilitate nuclear envelope rupture, followed by chromosome congression driven by the collapse of the nuclear F-actin network and capture by microtubules. Finally, the forming spindle is transported to the cell periphery by cytoplasmic flows produced by a wave of cortical contraction. Together, the presence of these actin-driven mechanisms in a basal metazoan evidence their ancient origin, establishing a metazoan complement of conserved molecular modules required for oocyte divisions.

## Introduction

Oocyte meiosis is a specialized form of cell division adapted to produce a fertilizable egg. It differs from somatic cell mitosis in important aspects; firstly, meiosis is a reductional division that produces a haploid egg. Therefore, the meiotic cell cycle is adapted to undergo two divisions in rapid succession without DNA replication. Secondly, oocytes are very large, storing nutrients to support early development of the embryo. To avoid dividing up these nutrients accumulated during oogenesis, oocyte divisions are extremely asymmetric. Of the four haploid sets of genome produced in meiosis, three are extruded into tiny, unviable polar bodies, resulting in a single, large haploid egg retaining all the cytoplasm and stored yolk, RNA and proteins within.

These oocyte-specific functions require substantial adaptations of the cytoskeletal systems. In somatic cell mitosis, dynamic centrosome-nucleated microtubule asters efficiently search and quickly capture chromosomes^1–3^. By contrast, oocytes have very large nuclei (the germinal vesicle) with a diameter of typically 50, in some species up to even 500 μm^4^. This constitutes a major challenge because, as predicted by theory^5^ and demonstrated experimentally^1,2,6^, microtubule ‘search and capture’ becomes inefficient in cells larger than the ∼30 μm diameter of somatic cells. Therefore, oocyte meiosis requires adapted mechanisms for efficient chromosome capture. Further exacerbating this, oocytes typically lack centrosomal microtubule asters, as centrosomes need to be eliminated during oogenesis in preparation for fertilization^7^. Even in species in which centrosomes are retained, they are inactivated to minimize the size of the polar body^8^. As a result, across metazoa, meiotic spindles are anastral and are much smaller than somatic spindles^9^. This necessitates adapted mechanisms not only for chromosome capture, but also for positioning the spindle and chromosomes within the oocyte. Yet another challenge is that the enormous oocyte nucleus features a specialized nuclear envelope packed with nuclear pore complexes and stabilized by a thick lamina^10^. In somatic cells, rapid disassembly and removal of the nuclear membrane is essential to allow access and enable efficient capture of chromosomes by cytoplasmic microtubules^11,12^. Therefore, oocytes further require specialized mechanisms to facilitate disassembly of the nuclear envelope.

Actin-driven mechanisms play a major role in mediating these oocyte-specific functions ‘helping out’ microtubules -- which primarily drive these processes in somatic mitosis. Prominently, actin-driven mechanisms have been shown to facilitate chromosome congression in oocytes of various species. In starfish, a contractile actin filament (F-actin) network transports chromosomes to cortically positioned microtubules, promoting their efficient capture^13–15^. More recently, Harasimov and coworkers showed that, strikingly similar to starfish, in human and porcine oocytes F-actin cables cluster chromosomes, essential to prevent chromosome loss^16^. While detailed mechanisms are less clear, *Xenopus* oocytes also feature an extensive nuclear F-actin network^17^, and reorganization of this network is essential for congression and transport of chromosomes to the oocyte cortex^18–20^.

Once chromosomes are captured and the spindle starts to assemble, F-actin has additional functions. In mouse oocytes, the spindle is transported from the center to the cortex by an actin-driven mechanism^21–23^. The mechanisms of this transport have been studied in substantial detail, revealing a dynamic cytoplasmic F-actin network nucleated by Formin-2 and Spire1/2, through which transport is driven by Myosin-5b and Myosin-2 motors^24,25^. The cytoplasmic network is functionally coupled to cortical F-actin nucleated by the Arp2/3 complex, whereby Arp2/3-mediated thickening of the cortex leads to the release of Myosin-2 to the cytoplasm^26,27^. In addition, cytoplasmic flows have been shown to contribute to transporting the spindle towards the cortex^28,29^. While the exact mechanisms are still unclear, Arp2/3-mediated filament nucleation is thought to generate a flow outwards of the ‘actin cap’ forming above the spindle, circulating back at the oocyte center, pushing the spindle toward the cortex^28,29^.

In addition, an ‘F-actin spindle’ has also been documented in mouse oocytes; a dense network of long actin filaments interwoven with spindle microtubules that facilitate the faithful segregation of chromosomes^30^.

Yet another, unexpected F-actin-driven mechanism has been observed early in meiotic maturation, so far only in starfish oocytes. At nuclear envelope breakdown (NEBD), a transient, Arp2/3-nucleated F-actin shell assembles on the nuclear side of the nuclear envelope that functions to rupture the nuclear envelope by prying apart nuclear membranes and the nuclear lamina^31,32^. This mechanism is necessary to dismantle the exceptionally robust nuclear envelope of oocytes, allowing cytoplasmic microtubules access to chromosomes^31,32^.

Finally, in the large oocyte, actomyosin-driven cortical contractions frequently form spatial waves moving across the oocyte, typically coinciding with cell cycle transitions^33,34^. These contractions are triggered by activation of the highly conserved RhoA pathway, causing surface contraction waves that drive large-scale cortical deformations and resulting cytoplasmic flows^35,36^, with functions still largely unclear.

Taken together, ample evidence from diverse species shows that the actin cytoskeleton mediates essential functions supporting the specialized divisions of oocytes; it facilitates nuclear envelope rupture, transports chromosomes, positions the spindle, and ensures faithful chromosome segregation. A critical, so far unaddressed question is whether these represent individual adaptations that emerged independently in certain species or clades, or whether they are broadly conserved mechanisms common to metazoan oocytes. Indeed, both scenarios are plausible: oocytes are one of the most diverse cell types showing an enormous variability in size and variable morphologies^10^. This reflects the adaptation of species to diverse environments, requiring diverse reproductive strategies. Thus, it would not be unexpected to find adaptations that are specific to oocytes of individual species or clades. On the other hand, there are indications for conserved mechanisms common to animal oocytes. This is reflected by shared features of overall morphology and cellular mechanisms, as well as molecular markers common to metazoan oocytes. A prominent example is the Mos kinase that is expressed exclusively in oocytes across metazoan species, and mediates key meiosis-specific functions such as the meiosis-specific cell cycle, and meiosis-specific spindle morphology^37,38^.

To address this question, we investigated oocyte meiosis in the non-bilaterian cnidarian jellyfish, *Clytia hemisphaerica*. Cnidarians are at the base of metazoan phylogeny and diverged from the bilaterian lineage more than 500 million years ago^39^. Thus, as an outgroup to Bilateria, *Clytia* is an ideal model for examining the evolutionary conservation and diversification of cellular mechanisms. Importantly, and unique among cnidarian model species, *Clytia* offers optically clear oocytes and hormonal control of meiotic maturation, allowing meiotic events to be analyzed by high-resolution live and fixed-cell confocal microscopy^40^. Indeed, conservation of the above-mentioned Mos kinase was demonstrated in *Clytia*, exemplifying this research strategy^41^.

Here, we first established fluorescent markers and quantitative live-cell imaging assays to visualize cytoskeletal dynamics in *Clytia* oocytes. Combining these with molecular perturbations and biophysical modeling revealed essential functions of actin through stages of meiosis. Upon meiotic entry, concomitant with NEBD, we observed a prominent F-actin shell forming on the nuclear side of the nuclear envelope, and could show that it facilitates nuclear envelope rupture. Secondly, we visualized an extensive F-actin network in the oocyte nucleus, and show that disassembly of this network aids chromosome congression. Thirdly, we show that the forming spindle is transported to the cortex by cytoplasmic flows produced by cortical contractions. Taken together, these findings evidence conserved roles of the actin cytoskeleton in oocyte meiosis in metazoa, defining core functional modules supporting the specialized oocyte divisions.

## Results

### High-resolution imaging of meiotic maturation in live Clytia oocytes

Our work was motivated by seminal contributions of the Houliston and Momose laboratories demonstrating how a cnidarian species, as an outgroup to Bilateria, can reveal conservation of fundamental cellular mechanisms of metazoan oocyte and embryo development^40,41^ (Fig. 1A).

**Figure 1.**
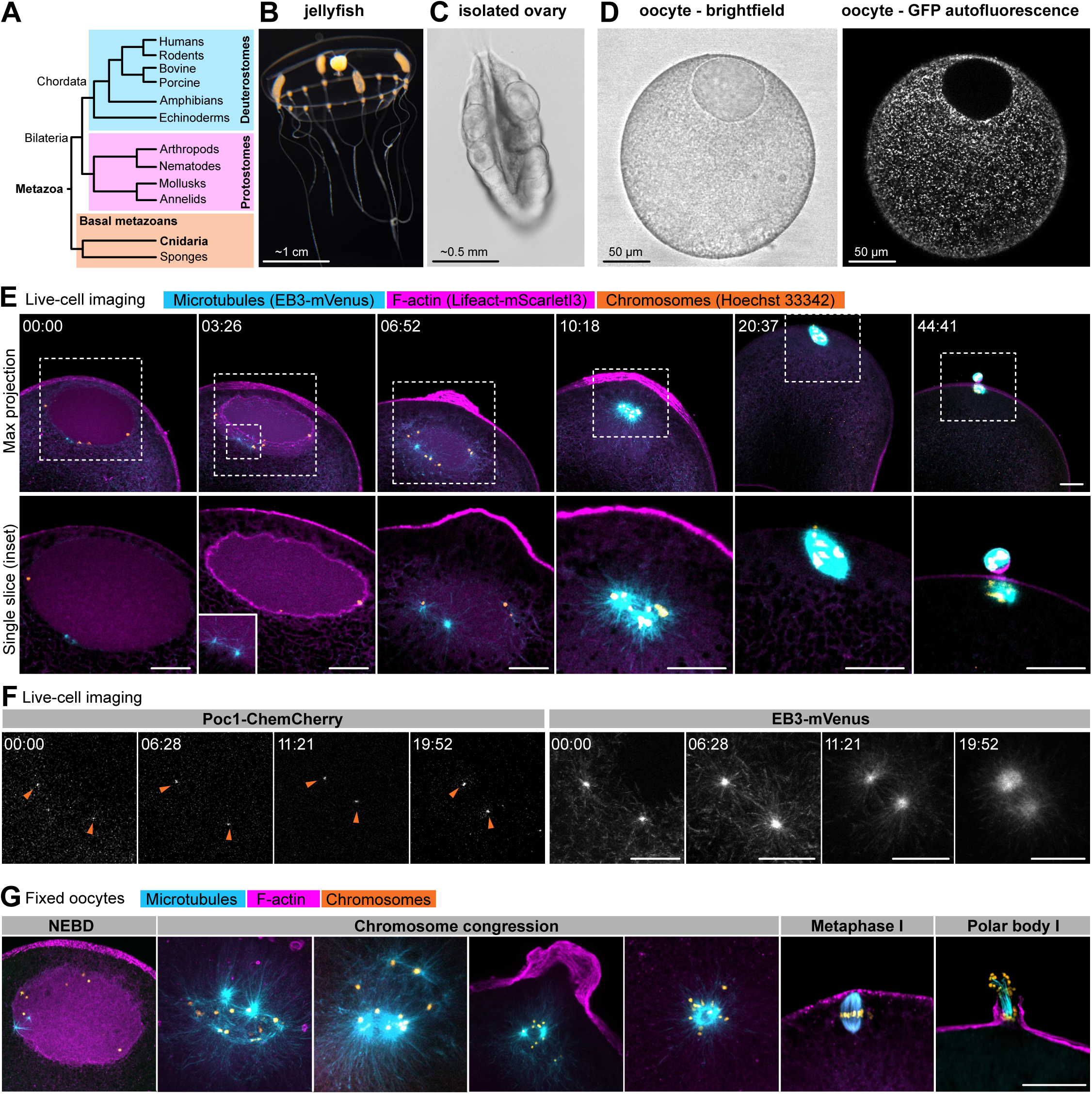
Imaging cytoskeletal dynamics during meiosis in *Clytia hemisphaerica* oocytes. **(A)** Simplified phylogenetic tree illustrating the relation of Cnidarian to major metazoan clades, highlighting those groups used for oocyte research. **(B)** Photograph of an adult *Clytia hemisphaerica* jellyfish. **(C)** Isolated ovary containing oocytes at different stages of development. **(D)** Isolated fully grown oocyte showing the cortically positioned nucleus by transmitted-light imaging (left), and endogenous GFP fluorescence in mitochondria (right). **(E)** Top panel: Selected maximum-intensity z-projections from a 3D confocal time series of the nuclear region of an oocyte. Actin is labeled with LifeAct-mScar-letI3 (magenta), microtubule plus-ends with EB3-mVenus (cyan) and chromosomes with Hoechst 33342 (orange). Lower panel: single confocal sections of the enlarged view of the regions indicated by dashed boxes. Scale bars: 20 μm. **(F)** Selected maximum-intensity z-projections from a confocal time series showing Poc1-ChemCherry (left) and EB3-mVenus (right) during meiotic maturation. Orange arrowheads indicate Poc1-ChemCherry-labeled centrioles. Scale bars: 10 µm. Time is shown as min:s. **(G)** Maximum-intensity projections of fixed oocytes from nuclear envelope breakdown (NEBD) through chromosome congression, metaphase I, and first polar body extrusion. Oocytes were stained for tubulin (cyan), F-actin (magenta), and chromosomes (orange). Scale bar: 20 μm.

We followed their detailed culture protocols to raise adult medusae measuring 2-3 cm in diameter with 4 radially arranged ovaries, containing oocytes at various stages of development^42^ (Fig. 1B, C). We manually isolated fully grown oocytes (∼170 μm diameter) showing the overall morphology of a typical metazoan oocyte with the nucleus located asymmetrically at the cortex, defining the animal-vegetal axis (Fig. 1D). As a peculiar feature, *Clytia* oocytes endogenously express a bright, mitochondria-targeted green fluorescent protein (GFP) (Fig. 1D).

To visualize cytoskeletal dynamics during meiotic divisions, we purified a set of heterologous recombinant protein markers, including the actin-binding peptide LifeAct^43^ fused to mScarletI3 (LifeAct-mScarletI3), and the microtubule plus-end binding protein EB3^44^ fused to mVenus (EB3-mVenus). Fortunately, at 514 nm excitation, mVenus is spectrally well separated from the endogenous GFP, allowing simultaneous two-color imaging with mVenus and mScarletI3. For staining chromosomes, we either used the cell-permeable dye Hoechst 33342 as a third label, or injected an anti-histone nanobody^45^ fused to mVenus or mScarletI3.

After injection of these markers, an accumulation of LifeAct in nuclei of immature oocytes was evident. In live oocytes, we could just barely resolve filaments (Fig. 1E), while in fixed samples, a nuclear F-actin network was readily visualized (Fig. 1G, and see below). Microtubules were abundant in the cytoplasm of immature oocytes, and showed no obvious organization; asters or nucleation centers were not distinguishable (Fig. 1E). Chromosomes were condensed and distributed along the periphery of the nucleus, likely kept in place by the nuclear F-actin network as demonstrated in *Xenopus* oocytes^46^.

Next, we induced meiosis by addition of the maturation hormone, WPRPamide. Within a few minutes, concomitant with NEBD, a prominent, transient F-actin structure assembled along the nuclear envelope. This F-actin shell then completely disassembled in 1-2 minutes -- very reminiscent of the F-actin shell observed in starfish oocytes facilitating nuclear envelope rupture^32^ (Fig. 1E).

At the same time, two microtubule asters emerged in close proximity to the nuclear envelope, typically on the vegetal side of the nucleus (Fig. 1E). The centriolar marker Poc1-ChemCherry^47^ localized to the centers of the two asters through meiosis (Fig. 1F). Poc1 localization and the constant number of two asters together indicate that *Clytia* oocytes do contain centrosomes, which are activated at induction of meiotic maturation. This is in contrast to acentrosomal microtubule organizing centers (aMTOCs), found in other species, for example in mouse oocytes forming in variable numbers^48^.

After NEBD, the microtubule asters continued to grow and moved into the area of the former nucleus (Fig. 1E, G). This was accompanied by the dissolution of the LifeAct signal in the nuclear region indicative of gradual disassembly of the nuclear F-actin network. Possibly facilitated by disassembly-mediated contraction of the F-actin network documented in starfish oocytes^15^, chromosomes were then captured by microtubule asters in less than 10 minutes (Fig. 1E, G). Next, the assembling spindle migrated towards the cortex, where it bipolarized (Fig. 1E, G). The bipolar spindle then rotated, followed by the extrusion of the first polar body, concluding the first meiotic division (Fig. 1E, G).

This entire process was accompanied by extensive changes in cortical F-actin dynamics driving rather dramatic shape changes. First, soon after NEBD, F-actin accumulated on the cortex at the animal pole forming a ‘bun-like’ protrusion, which then flattened out again by the time chromosomes were captured by microtubules (Fig. 1E, G). This was then followed by a massive wave of cortical contraction culminating in polar body extrusion (Fig. 1E, and see below), reminiscent of the surface contraction waves seen for example in *Xenopus* or starfish oocytes^34,35^.

Together, by developing recombinant protein markers and high-resolution confocal microscopy, we were able to visualize meiotic events in *Clytia* oocytes, revealing prominent F-actin structures in immature oocytes, at NEBD, during chromosome transport, as well as changes in cortical dynamics.

### A nuclear F-actin network is present in immature Clytia oocytes

To explore when the nuclear F-actin network forms during oogenesis and to characterize its structure in more detail, we performed phalloidin stainings of whole ovaries, co-stained with the universal nuclear pore antibody mAb414 (Fig. 2A). This revealed a prominent nuclear F-actin staining at all stages of oocyte development, whereas somatic nuclei of endothelial cells only showed staining of cytoplasmic F-actin structures (Fig. 2A). A nuclear F-actin network was evident already in Stage I oocytes and was maintained across all stages of oogenesis (Fig. 2B). At later stages, the F-actin network developed to finer and denser with an approximate pore size around 1.5 μm in fully grown immature oocytes (Fig. 2C).

**Figure 2.**
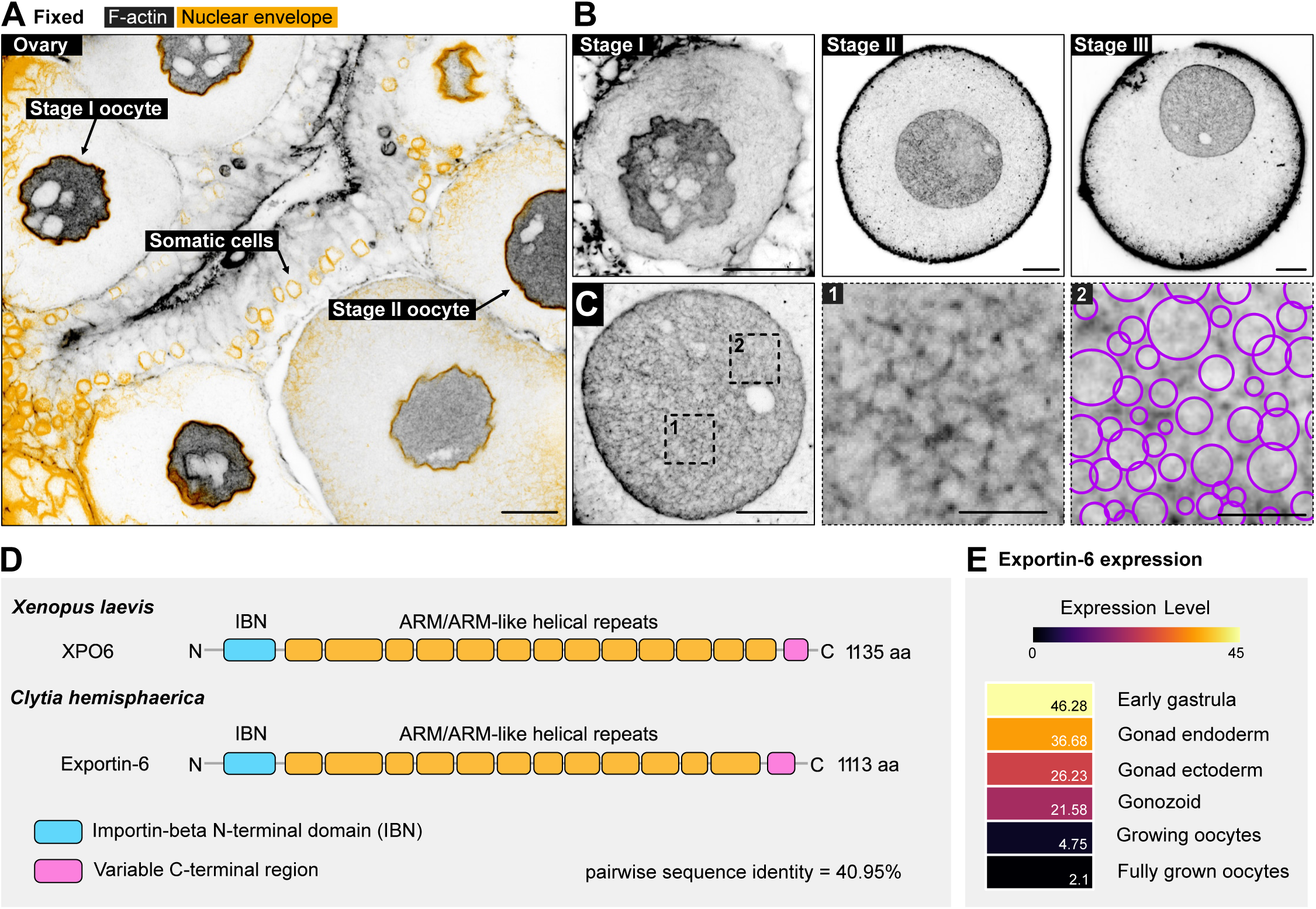
A nuclear F-actin network is present in immature oocytes. **(A)** Confocal images showing F-actin organization in fixed *Clytia* immature oocytes. Left: Isolated ovary stained for F-actin (gray) and the nuclear envelope (orange). Black arrows indicate stage I, stage II oocytes and endodermal somatic cells. Scale bars: 20 μm. **(B)** Stage I, stage II, and stage III oocytes showing F-actin (gray) organization during oocyte growth. **(C)** Stage III oocyte nucleus showing an F-actin network, with two regions indicated by dashed boxes shown in the zoom. Inset 2 is overlaid with purple circles indicating detected meshes for mesh-size quantification. Scale bars: 20 μm; insets: 10 μm. **(D)** Schematic illustration of Exportin-6 domain organization in *Xenopus laevis* and *Clytia hemisphaerica*. **(E)** Expression profile of the Exportin-6 in different cell types and tissues, with lowest expression observed in fully grown oocytes.

Across animal species, somatic nuclei typically do not contain such an extensive F-actin network or filamentous actin in general, because abundantly expressed Exportin-6 efficiently exports profilin-actin complexes from the nucleus^49^. However, in *Xenopus* an extensive F-actin network was shown to stabilize the gigantic, 0.5 mm nucleus^17,46^. This is caused by specific downregulation of Exportin-6 (XPO6) expression in oocytes, while most other *Xenopus* cell types do express Exportin-6^17^. To test whether a similar mechanism may be at work in *Clytia*, we searched for homologs of Exportin-6 in publicly available genomic resources. We were able to identify a *Clytia* Exportin-6 homolog featuring all essential functional domains of the vertebrate protein (Fig. 2D). Intriguingly, cell-type-specific transcriptome data^50^ reveal that Exportin-6 expression is specifically suppressed in developing and fully grown oocytes, while it is abundantly expressed in other cell types (Fig. 2E). This suggests that, similar to *Xenopus* oocytes, oocyte-specific downregulation of Exportin-6 expression may cause nuclear accumulation of F-actin in *Clytia* oocytes.

### A transient F-actin shell facilitates nuclear envelope rupture

Next, we focused on the prominent assembly of F-actin along the nuclear envelope that appeared a few minutes after initiation of meiotic maturation, at around the time of NEBD, and disassembled again in just 1-2 minutes (Fig. 3A). This transient F-actin structure showed a remarkable similarity to the ‘F-actin shell’ we discovered earlier in starfish oocytes^31,32^. Therefore, we wanted to explore the extent of structural and functional conservation between the two species.

**Figure 3.**
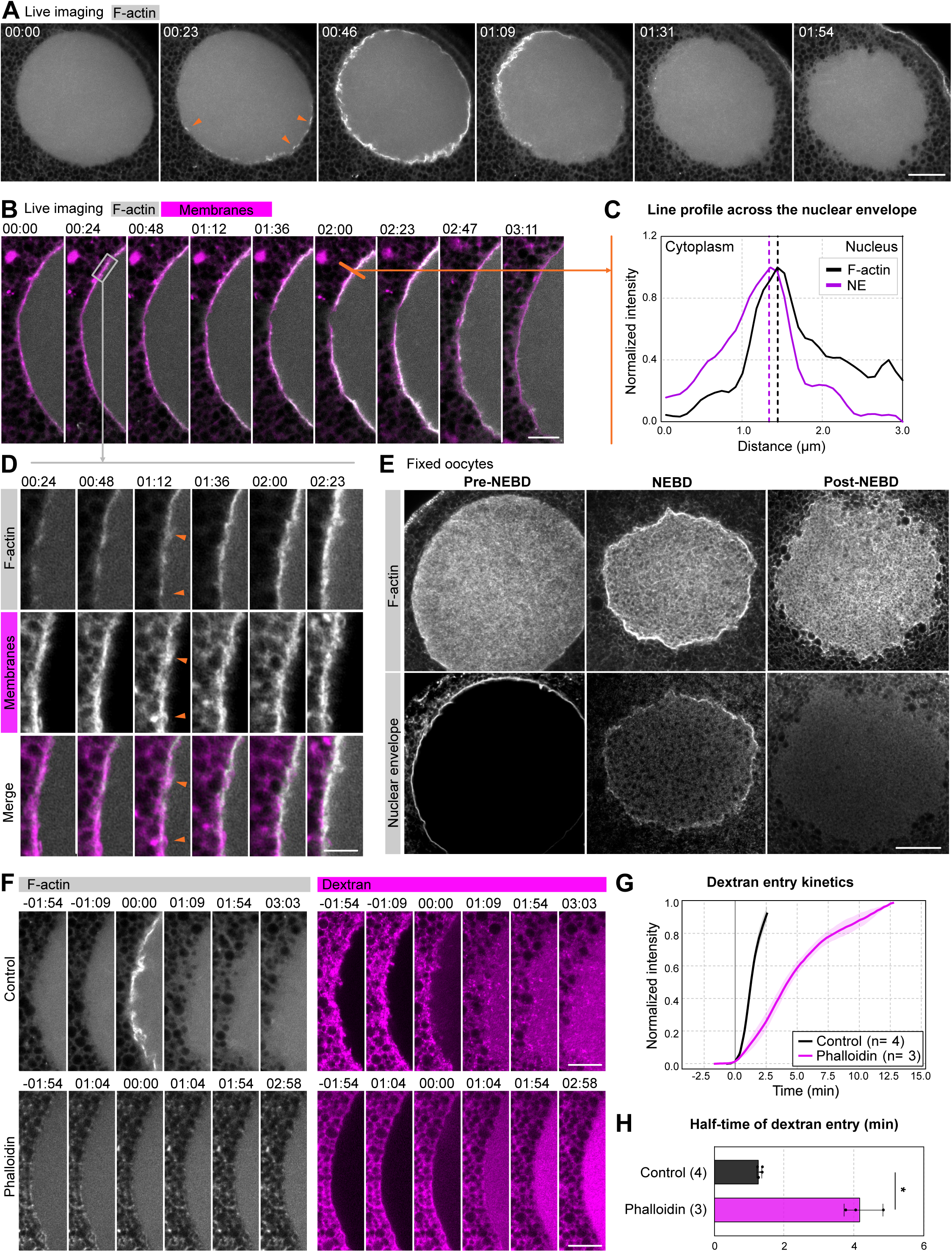
A transient F-actin shell forms underneath the nuclear envelope promoting its rupture. **(A)** Confocal sections of a live oocyte’s nuclear region showing formation and subsequent disassembly of the F-actin shell. Oocytes were injected with LifeAct-mScarletI3 to label F-actin (gray). Arrowheads indicate initial F-actin foci. Scale bar: 20 μm. **(B)** Confocal section of a live oocyte’s nuclear periphery, showing details of F-actin shell formation during NEBD. Oocytes were injected with LifeAct-mScarletI3 to label F-actin (gray) and DiIC_18_(5) to label endomembranes (magenta). The orange line indicates the position used for the line profile in (C). Scale bar: 20 μm. **(C)** Line profile of normalized intensity of F-actin (black) and the nuclear envelope (NE; purple) signal intensities. Dashed lines indicate the positions of peak F-actin and NE intensities, respectively. **(D)** Enlarged single-slice from the time series shown in (B) indicated by the boxed region. Arrowheads indicate local changes in F-actin and endo-membrane organization. Scale bar: 5 μm. **(E)** Confocal images of fixed oocytes stained for F-actin (left) and nuclear envelope (right) before, during, and after NEBD. Scale bar: 20 μm. **(F)** Confocal time-lapse series of the nuclear periphery in control and phalloidin-injected oocytes. Oocytes were co-injected with LifeAct-mScarletI3 to label F-actin (gray) and 500 kDa dextran (conjugated to Abberior Star Red) as a cytoplasmic probe, showing delayed entry in phalloidin treated oocytes. Scale bars: 10 μm. **(G)** Dextran entry kinetics in control and phalloidin-treated oocytes. Normalized dextran intensity obtained from the central nuclear region. The number of oocytes is indicated by n. Dashed line indicates NEBD. **(H)** Half-time dextran entry in control and phalloidin-treated oocytes. Individual points represent individual oocytes. Bars show [mean ± SEM]. Welch’s t test, p = 0.0113. Time is shown as min:s.

High-resolution live imaging of *Clytia* oocytes injected with LifeAct-mScarletI3 showed that, similar to starfish oocytes, the F-actin shell initiates at foci along the nuclear envelope, from which shell assembly spreads out in a wave-like manner (Fig. 3A, B). Simultaneous imaging of F-actin and endomembranes revealed that F-actin assembles on the nuclear side, and that the assembly of the F-actin shell coincides with the fragmentation of the nuclear membranes (Fig. 3B-D).

We confirmed the presence of these structures in fixed oocytes stained with phalloidin and mAb414 to visualize the nuclear envelope (Fig. 3E): prior to NEBD, nuclear pore staining formed a continuous boundary, and the F-actin network homogeneously filled the nucleus. At NEBD, concomitant with fragmentation of the nuclear envelope, F-actin enriched along the nuclear periphery, while the F-actin network coarsened. After NEBD, nuclear pore staining progressively diminished, while F-actin network signal persisted, suggesting that F-actin shell may facilitate nuclear envelope rupture.

To functionally assess whether the F-actin shell has a role in nuclear envelope rupture, as it is the case in starfish oocytes, we next monitored the entry kinetics of a cytoplasmically injected fluorescent 500-kDa dextran into the nucleus^51,52^. In control oocytes, dextran entry correlated closely with F-actin shell formation and rapidly filled the nucleus (Fig. 3F). Next, we depleted actin monomers by injecting a large amount of phalloidin, which drives all available actin into stable filaments, and thereby prevents formation of any new F-actin structure^32^. Indeed, phalloidin injection effectively prevented the formation of the F-actin shell (Fig. 3F). Additionally, phalloidin injection significantly delayed dextran entry (Fig. 3G). The half-time of dextran entry increased about 4-fold from ∼1 min in control oocytes to ∼4 min in phalloidin-injected oocytes, evidencing that the F-actin shell facilitates permeabilization of the nuclear envelope (Fig. 3H).

Collectively, our data reveal a remarkable structural and functional conservation of the F-actin shell between starfish and *Clytia* oocytes. In both species, the F-actin shell is initiated from foci, assembles on the nuclear side of the nuclear envelope, extends in a wave-like manner, and then disassembles again in 1-2 minutes. Both in *Clytia* and starfish, preventing F-actin shell assembly delays nuclear envelope permeabilization, as monitored by the kinetics of dextran entry. Together, these data evidence that the F-actin shell, documented previously in starfish oocytes alone, is not an adaptation specific to echinoderms but is a more broadly conserved feature of metazoan oocytes.

### Chromosome tracking separates two phases: congression followed by transport to the cortex

After the nuclear envelope and the F-actin shell disassemble, chromosomes first collect near the center of the former nucleus and are subsequently transported to the oocyte cortex (Fig. 4A). Next, we wanted to quantitatively characterize this ∼10-minute process by tracking chromosomes in 3D at high temporal resolution (∼5 s per volume). For this we developed a workflow to segment chromosomes and the surface of the oocyte and to automatically track chromosome motion (Fig. 4C). To characterize the congression phase, we determined the centroid (i.e. the mean of all) of chromosome positions, and calculated the intensity-weighted mean distance of chromosomes to the centroid at each time point (Fig. 4D, E). To quantify the transport of the chromosomes to the cortex, we calculated the nearest distance between the centroid and the cortex (Fig. 4D, F).

**Figure 4.**
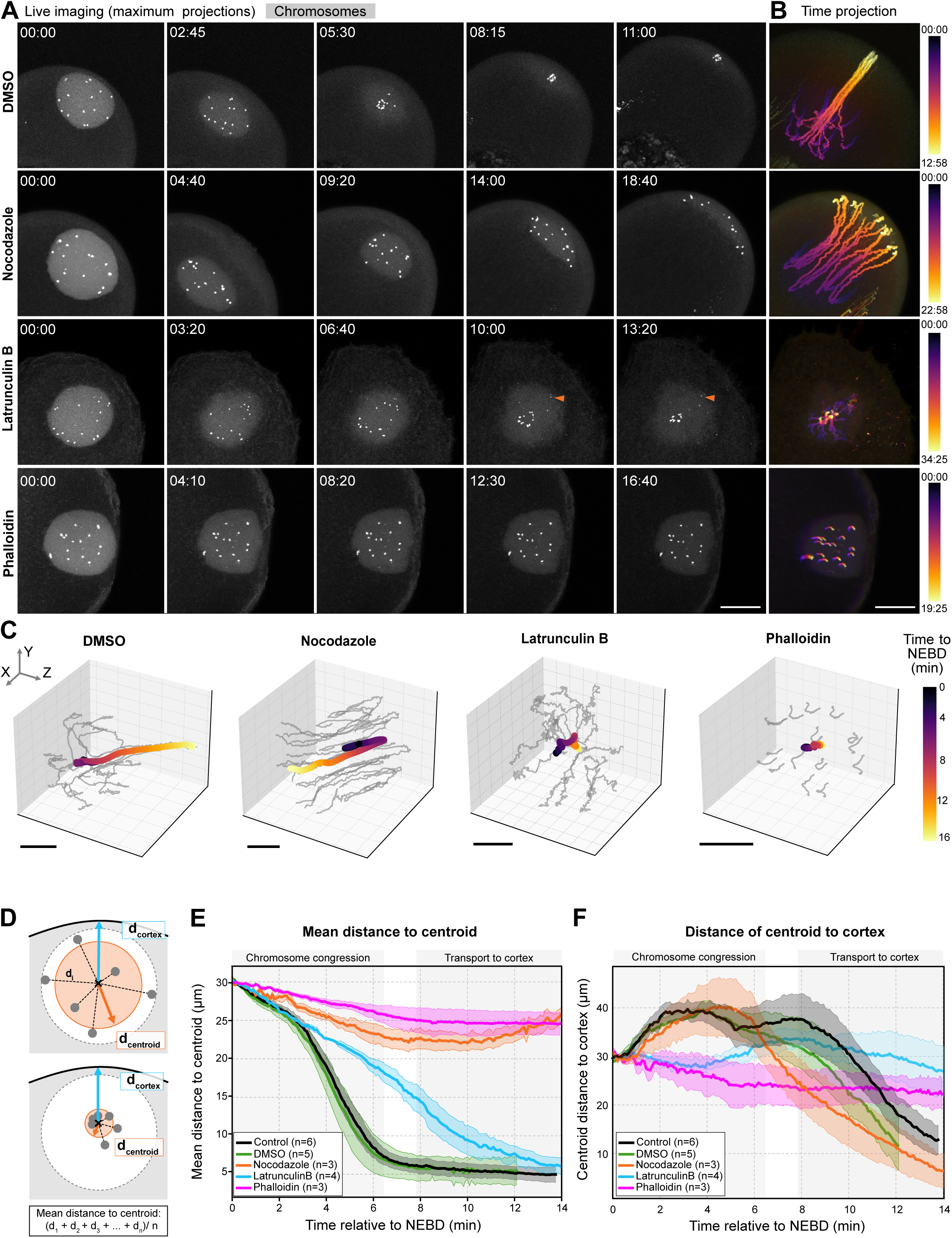
Actin and microtubules cooperate to collect and transport chromosomes. **(A)** Maximum-intensity projections from a 3D time-lapse series of a live oocyte’s nuclear region during chromosome congression and transport. Oocytes were injected with chromobody-mScarletI3 or chromobody-mVenus to visualize chromosomes (gray). Oocytes were treated with DMSO, nocodazole, latrunculin B and phalloidin as indicated. Arrowheads indicate lost chromosomes. Time is shown as min:s relative to NEBD. Scale bar: 50 μm . **(B)** Time projections of chromosome trajectories from the oocytes shown in (A). The pseudo color scale bar indicates time relative to NEBD. Scale bar: 50 μm. **(C)** 3D chromosome trajectories in DMSO-, nocodazole-, phalloidin- and latrunculin B-treated oocytes. Gray lines represent a single chromosome trajectories and colored lines represent the centroid of all chromosome positions over time. Scale bars: 25 μm. **(D)** Scheme illustrating the measurements used to quantify chromosome congression and transport. Chromosome congression was quantified as the mean distance of individual chromosomes to their centroid (d_centriod_, orange line) at each time point. The distance to the cortex (d_cortex_, blue line) was quantified as the shortest distance of centroid (d_centroid_) to the oocyte cortex. **(E)** Plot of mean chromosome distance to the centroid (d_centroid_) over time in control, DMSO-, nocodazole-, latrunculin B- and phalloidin-treated oocytes. **(F)** Plot of centroid distance to the oocyte cortex over time for the conditions shown in (E). In both (E) and (F) plots, line indicates mean values of multiple oocytes per condition and shaded region indicates SEM.

Next, to separate out contributions of the actin and microtubule cytoskeleton, we imaged oocytes in which we either: (i) depolymerized microtubules by nocodazole, (ii) depolymerized actin filaments by treating oocytes with latrunculin B, or (iii) stabilized actin filaments by injection of phalloidin. Neither of the treatments interfered with the initial timing of maturation and NEBD, while after NEBD the effects of the treatments were evident.

In control or DMSO-treated oocytes, chromosomes congressed into a compact cluster within ∼6 min after NEBD (Fig. 4A-E). The congression process was highly reproducible between oocytes, with an initial slower phase lasting 2-3 minutes, followed by a faster phase. By contrast, chromosomes in nocodazole-treated oocytes initially moved toward each other but then failed to cluster and remained dispersed in the nuclear region (Fig. 4A-E). In latrunculin B-treated oocytes, chromosomes did cluster, although with a slight delay. Chromosomes located near the nuclear periphery were frequently left behind and were captured only very late or not at all (Fig. 4A). Stabilizing actin filaments by phalloidin injection completely prevented chromosome congression (Fig. 4A, C).

The transport phase showed a very distinct dependency on cytoskeletal perturbations. In nocodazole-treated oocytes, despite failing to congress, individual chromosomes were efficiently transported to the cortex (Fig. 4A-F). By contrast, interfering with the actin cytoskeleton blocked chromosome transport; both phalloidin stabilization and latrunculin B treatment prevented chromosome movement to the oocyte cortex (Fig. 4A-F).

Together, these data indicate that the first congression phase is primarily driven by microtubules but also critically depends on F-actin disassembly, as filament stabilization prevents congression, while accelerated disassembly leads to chromosome loss. The subsequent transport of the chromosomes to the cortex is independent of microtubules and strictly depends on an intact actin cytoskeleton.

### Combined F-actin network disassembly and microtubule capture explains chromosome congression

We first focused on the mechanism of the congression phase. During this phase, we were able to image chromosome and microtubule dynamics at high resolution and in 3D (Fig. 5A), however, we had insufficient resolution to visualize the F-actin network and its disassembly dynamics in live oocytes. Therefore, we used computer simulations to recapitulate the observable morphology of the cytoskeletal elements and 3D chromosome trajectories, and thereby identify potential mechanisms.

**Figure 5.**
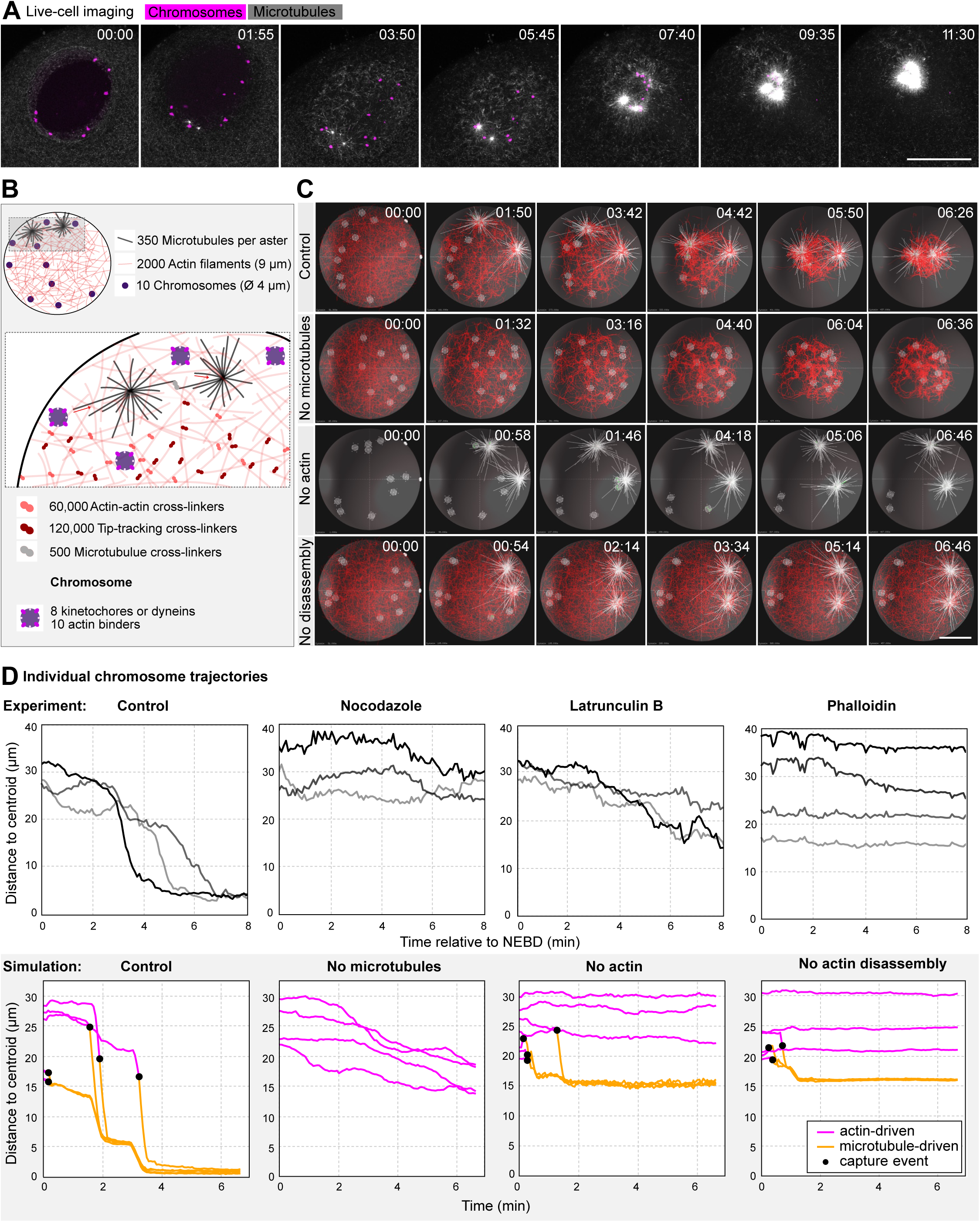
Transport by F-actin network and microtubule capture explains chromosome congression. **(A)** Selected maximum-intensity projections from a confocal series of chromosome congression in live oocytes. Oocytes were injected with chromobody-mScarletI3 to label chromosomes (magenta) and with EB3-mVenus to label microtubule plus ends (gray). Scale bar: 50 μm. Time is shown as min:s. **(B)** Schematic illustration of 3D Cytosim simulation of chromosomes congression. Chromosomes are represented as spherical objects (purple) which interact with the F-actin network (red) and microtubules (gray) through actin-binders and kinetochores, respectively. The indicated numbers and dimensions correspond to the parameters used in the simulation. **(C)** Renderings of 3D Cytosim simulations of chromosome congression under control conditions, in the absence of microtubules, in the absence of actin and when actin disassembly is prevented. Microtubules are shown in gray, actin filaments in red and chromosomes are gray spheres with kinetochores. Scale bar: 20 μm. Time is shown relative to NEBD in min:s. **(D)** Top: Plot of the distance to centroid of selected individual chromosome trajectories from control, nocodazole-, phalloidin- and latrunculin B-treated oocytes. Bottom: Plot of distance to centroid of selected individual chromosome trajectories from simulations shown in (C). Actin-driven transport is represented as magenta lines and microtubule driven phase as orange lines, black dots indicate chromosome capture events.

Given the overall similarity of the process to chromosome congression in starfish oocytes, we used models we previously implemented in the Cytosim environment for starfish oocytes as a starting point^53^. In these works, we separately modeled chromosome transport mediated by a disassembling F-actin network^15^, and chromosome capture by microtubules^54^. Here, we combined these two models preserving most previously validated parameters, and made the minimal necessary modifications to adjust for *Clytia*-specific features (Table S1).

For the F-actin network, we implemented the same disassembly-driven contractile mechanism as in starfish oocytes that is based on cross-linkers tracking the depolymerizing filament end (Fig. 5B). Specifically, actin filaments were connected both by tip-tracking and filament-to-filament cross-linkers. Matching experimental observations, we started the simulation with a pre-assembled network and set the parameters such that the network disassembles in approximately 6 min, by the end of the congression process. We also extended the previous 2D simulation, to which we were constrained to due to limited computational resources at the time, to a 3D simulation of the F-actin network. Regarding microtubules, we changed the model such that microtubule asters are not anchored to the cell cortex, as they do in starfish oocytes. Instead, asters are initially positioned at the nuclear envelope and can then move freely into the nuclear area after NEBD. Otherwise, microtubule dynamic instability was simulated using typical parameters (Fig. 5B). Chromosomes were simulated as simplified, spherical objects, each with 8 ‘kinetochores’, i.e. microtubule binding sites, which also have dynein activity transporting attached chromosomes poleward (Fig. 5B).

Simulations recapitulated key experimental observations. In the ‘control’ unperturbed condition, chromosomes embedded in the F-actin network were collected by the disassembling F-actin network, moving chromosomes closer to microtubule asters, facilitating their capture (Fig. 5C). When comparing individual chromosome trajectories from experiments and simulations, velocities and overall temporal dynamics were similar; as in experiments, we could distinguish an initial, slower, actin-driven phase followed by capture and fast pole-ward transport on microtubules (Fig. 5D).

In simulations without microtubules, the F-actin network moved chromosomes somewhat closer to one another, but they remained scattered, as observed in nocodazole-treated oocytes (Fig. 5C). If actin filaments were removed, nearby chromosomes were efficiently captured by microtubule asters, however chromosomes distal to microtubules were left behind, just as observed in latrunculin B-treated oocytes (Fig. 5C, D). Stabilization of the filament network impaired chromosome capture, consistent with observations in phalloidin-injected oocytes. However, in simulations, chromosomes located in close proximity were still captured by microtubules, whereas in experiments chromosome capture was completely blocked. This discrepancy likely results from the lower F-actin network density we were able to simulate as compared to the experimental system (Fig. 5C).

Taken together, our simulations recapitulate chromosome capture by microtubule asters facilitated by a disassembling F-actin network, that together constitute a robust system for chromosome congression in large nuclei of oocytes. Simulations highlight the necessary balance between F-actin network stability and disassembly; disassembly drives contraction, and it is also required to release chromosomes for microtubule capture. However, if F-actin network disassembly is too fast, nearby chromosomes will be captured faster, but distal chromosomes are lost as the F-actin network loses connectivity -- matching observations in latrunculin B-treated oocytes. Slow disassembly, on the other hand, holds chromosomes back, hindering capture by microtubules recapitulating our observation in phalloidin injected oocytes.

### Cytoplasmic flows transport chromosomes to the cortex

After congression at the center of the former nucleus, chromosomes are transported to the cortex (Fig. 4A). This step lasts ∼5 minutes and coincides with cytoplasmic flows in the bulk, well-visible by transmitted light imaging, and even clearer by imaging of the endogenous GFP fluorescence in mitochondria (Fig. 6A). Based on this correlation, we hypothesized that cytoplasmic flows may contribute to chromosome transport. As a first test, we used particle image velocimetry (PIV) to quantify flow fields, and compared chromosome velocities derived from chromosome tracking with local PIV flow velocities (Fig. 6B).

**Figure 6.**
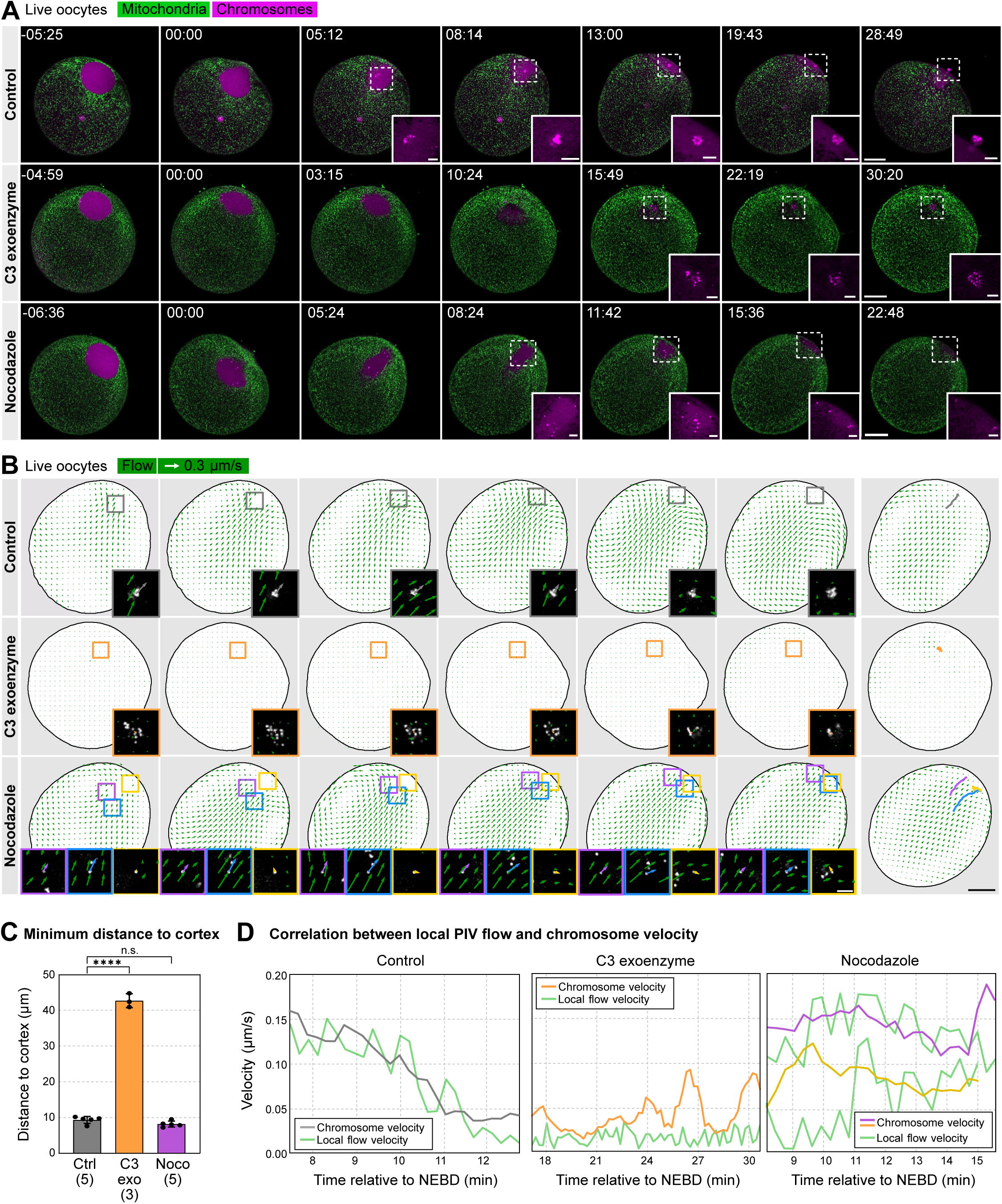
Cytoplasmic flow transports congressed chromosomes to the cortex. **(A)** Confocal slices selected from a time lapse series of live oocytes showing cytoplasmic mitochondria, visualized by endoGFP (green), and chromosomes (magenta, Chromobody-mScarletI3) in control, C3 exoenzyme- and nocodazole-treated oocytes. Insets show the chromosome channel in the regions marked by dashed squares. Time is given as min:s. Scale bars: 50 μm, insets: 10 μm. **(B)** Cytoplasmic flows analysed by particle image velocimetry (PIV) of mitochondrial endoGFP signal in the oocyte cytoplasm, for all conditions shown in (A). Green vectors indicate the direction and magnitude of the flow field. Insets correspond to the boxed region and show chromosomes overlaid with the local PIV vectors and chromosome velocity vector. The final image in each condition shows chromosome trajectories over time for clustered chromosomes in control (gray) and C3 exoenzyme (orange) and for nocodazole in purple, blue and yellow for individual chromosomes. Scale bar: 50 μm, insets: 10 μm. **(C)** Minimum distance of chromosomes to the oocyte cortex in control, C3 exoenzyme- and noco-dazole-treated oocytes. n is indicated in brackets. One-way ANOVA test; *p <0.0001*. **(D)** Plots showing correlation between the magnitude of chromosome velocity and local PIV flow velocity during chromosome transport to cortex in control, C3 exoenzyme- and nocodazole-treated oocytes. Local flow velocity was obtained from the PIV vectors surrounding the corresponding chromosome. Correlation plots used the same time intervals shown in (B). Time is shown in min:s.

In control oocytes, the onset of the flow coincided with the movement of the cluster of congressed chromosomes toward the cortex at the animal pole. The velocity of the chromosome cluster closely correlated with the local flow velocity spatially (Fig. 6B) and temporally (Fig. 6D), suggesting that cytoplasmic flows may indeed transport chromosomes.

Next, we attempted to inhibit cytoplasmic flows using the C3-exoenzyme, a general inhibitor of the Rho-pathway and thus cellular contractility. Indeed, C3-exoenzyme treatment completely abolished cytoplasmic flows (Fig. 6A, B). In these oocytes, NEBD occurred with normal timing, and chromosomes were efficiently congressed, indicating that C3-exoenzyme does not interfere with cytoplasmic F-actin structures and microtubule dynamics up until this stage. By stark contrast, chromosomes failed to move toward the cortex and remained at the site of congression, suggesting that inhibition of cytoplasmic flows blocks chromosome transport (Fig. 6A-D).

We additionally analyzed chromosome transport in nocodazole-treated oocytes. As we show above, depolymerization of microtubules interferes with chromosome congression, but individual chromosomes are nevertheless transported to the cortex (Fig. 4A; 6A). Analysis of flow fields revealed that nocodazole treatment does not significantly affect cytoplasmic flows, evidencing that they are independent of microtubules (Fig. 6A, B). Strikingly, in nocodazole-treated oocytes, velocities of individual, scattered chromosomes correlated with local PIV flow velocities both spatially and temporally (Fig. 6A-D). This strongly indicates that chromosomes are transported by cytoplasmic flows, independently of whether they are clustered by microtubules or are individual objects.

Together, the tight spatiotemporal correlation combined with perturbation experiments evidence that transport of chromosomes to the oocyte cortex is driven by cytoplasmic flows. Nocodazole treatments further show that coupling occurs at the level of chromosomes, and is independent of microtubules.

### Cytoplasmic flows are driven by cortical contraction

The C3-exoenzyme is an effective inhibitor of the RhoA pathway, a key regulator of cortical contractility^35^. Thus, the observation that C3-exoenzyme injection inhibits cytoplasmic flows suggests that flows may be driven by cortical contractility. To explore this hypothesis, we first imaged cortical F-actin (Life-act-mScarletI3) as a proxy for cortical contractility (Fig. 7A). In the same oocytes, we also imaged the endogenous GFP to directly correlate changes in cortical F-actin to flow fields. This revealed a strong correlation between changes in cortical F-actin and cytoplasmic flows (Fig. 7A, B).

**Figure. 7.**
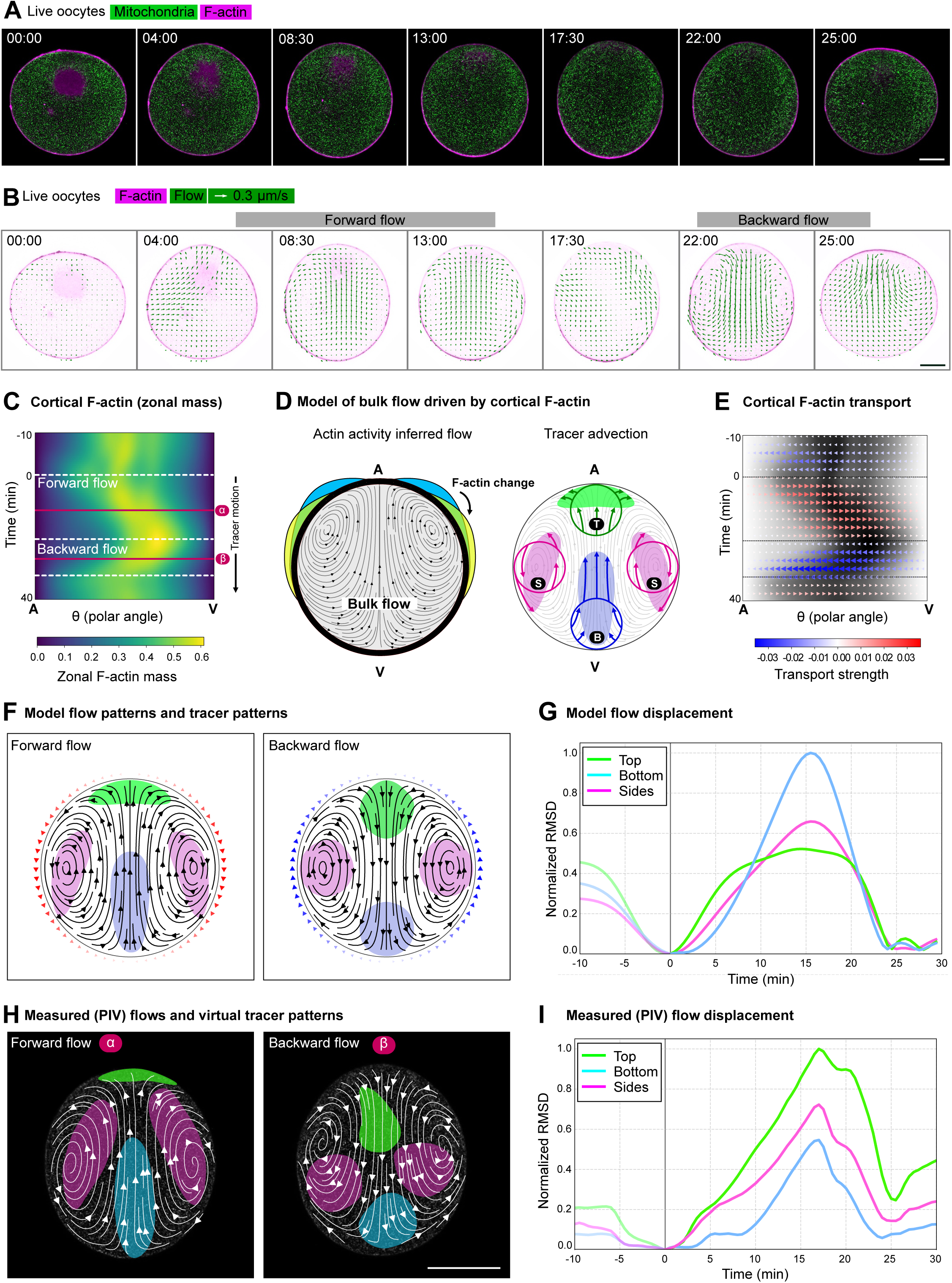
Bulk cytoplasmic flow is driven by cortical contraction. **(A)** Confocal slices selected from a time lapse series of live oocytes showing cytoplasmic mitochondria, visualized by endogenous GFP (green), and F-actin (magenta, LifeAct-mScarletI3). **(B)** Cytoplasmic flows analysed by PIV on the mitochondrial endogenous GFP. Green vectors indicate the direction and magnitude of the flow field. F-actin signal is in magenta as shown in (A). Scale bar: 50 μm. Time is shown in min:s. **(C)** Kymograph of the measured actin distribution (zonal F-actin mass) in the cortex. The polar angle is measured meridionally from the animal pole (A) to the vegetal pole (V). The two distinct flow phases are marked by dashed lines; the tracer simulation starts with the onset of the forward flow. **(D)** Biophysical model consisting of a spherical Stokes flow with no-slip boundary conditions on the outer shell in which the bulk flow is driven by tangential, meridional deformation of the outer membrane dragging the fluid with it. Left: boundary movement is extracted from the change in cortical F-actin intensity obtained from one frame to the next. Cortical F-actin motion drives 3D Stokes flow in the bulk of the spherical simulation domain. Right: virtual, passive tracers are simulated in the flow to analyze the cytoplasmic material deformation. Three distinct tracer regions are used near the animal pole (<u>T</u>op, green), the equator (<u>S</u>ides, magenta), and near the vegetal pole (<u>B</u>ottom, blue). The time-dependent flow advects the tracer regions. Depicted are the initial, circular starting locations of the tracers and during maximum deformation. **(E)** Reconstructed cortical F-actin motion by solving the transport equation from one frame to the next, depicted as meridional arrows. The resulting transport is used as the boundary condition for the bulk flow. The F-actin mass distribution is shown in grey. **(F)** Virtual passive tracer regions are advected in the experimental flows, for comparing the material deformations seen in the experiment and the simulation driven by the F-actin motion. **(G)** Mean tracer region displacement for reconstructed flows. **(H)** Experimental flow fields obtained from PIV, averaged over 5 min, marked with α and β on (C). Scale bar: 50 μm. **(I)** Mean tracer region displacement for experimental flows.

Analyzing the entire process identified two distinct phases: a forward flow, which coincides with chromosome transport, we focused on above, followed by a backward flow coinciding with the subsequent polar body formation (Fig. 7B). During the first phase, concomitant with the onset of the cytoplasmic flows, cortical F-actin began to develop an asymmetry. F-actin intensity decreased at the animal pole and enriched at the vegetal pole, moving as a wave across the oocyte (Fig. 7A, B). During the second phase, the wave reversed direction and moved backwards from the vegetal to the animal pole, correlated with the backward flow. This relocalization of cortical F-actin, first concentrating at the vegetal pole and then moving towards the animal pole is well visualized on a kymograph of the zonal F-actin mass (Fig. 7C).

To explain the observed correlation between cortical F-actin and flows, we implemented a biophysical model, whereby the bulk flow is driven by tangential, meridional deformation of the outer membrane, caused by cortical contraction (Fig. 7D). In the model, the deformation of the outer membrane is directly proportional to the change in cortical F-actin. Thus, the wave of change in F-actin intensity causes a wave for deformation, which drags the underlying cytoplasm with it, resulting in bulk flows (simulated as a spherical Stokes flow with no-slip boundary conditions on the outer shell) (Fig. 7D). To relate these flows to chromosome transport, we also simulated virtual tracers that are carried with the flow (Fig. 7D).

Therefore, we first extracted the boundary movement from the cortical F-actin intensity change (i.e. F-actin transport, Fig. 7E). We then simulated the 3D Stokes flow in the bulk in a spherical simulation domain. Strikingly, these simulations recapitulated flow patterns very similar to patterns measured by PIV analysis (compare Fig. 7F and 7H): the wave of cortical F-actin moving from the animal to the vegetal pole drove cortical flows in the same direction, which then caused a forward flow in the middle of the oocyte from the vegetal to the animal pole (Fig. 7F, H). The flow pattern then reversed when F-actin moved in the opposite direction in the second phase (Fig. 7F, H). As visualized by virtual tracers, during the forward flow the cytoplasmic material was transferred towards and pushed against the animal pole causing it to spread laterally (Fig. 7F, H). The backward flow then transferred the material back near to the initial position (Fig. 7F, H). The temporal kinetics is visualized by plotting the mean tracer region displacements over time both for simulation and experiment (Fig. 7G, 7I).

Displacement of virtual tracer positions were remarkably similar to chromosomes, particularly well visible in nocodazole-treated oocytes, in which chromosomes are scattered (see Fig. 6 and above). However, unlike tracers, chromosomes are not moved backwards in the second phase by the backward flow. This is likely explained by the fact that the microtubules of the assembling spindle anchor to the chromosome cluster to the cortex. Indeed, when microtubules are depolymerized, a backward movement of chromosomes is observed.

Taken together, our observations combined with biophysical modeling provide a simple explanation for how waves of cortical contractions can drive cytoplasmic flows in the bulk, driving transport of chromosomes to the oocyte cortex.

## Discussion

We provide here the first detailed characterization of cytoskeletal dynamics during oocyte meiosis in a cnidarian species, taking advantage of newly developed recombinant fluorescent markers and quantitative live cell imaging assays. We find that the overall morphology and progression of oocyte meiosis in *Clytia* is similar to that of other metazoan, bilaterian, and in particular to deuterostome species. A likely consequence of the similar marine habitat, *Clytia* oocytes are remarkably similar in size and morphology to starfish (*Patiria*) oocytes, which we extensively studied in earlier work. This provided us with a unique opportunity to compare the two, otherwise phylogenetically distal species diverged more than 600 million years ago^39^.

Our data revealed multiple actin-driven mechanisms supporting oocyte divisions in *Clytia*, showing striking similarity to mechanisms documented in other species. Thus, a key conclusion of our work is that actin-driven mechanisms are conserved features common to metazoan oocytes, evidenced by their presence in oocytes of an early branching non-bilaterian species. Specifically, we identify at least three core molecular modules adapted to meiosis-specific functions in oocytes: (i) an F-actin shell that facilitates the rupture of the nuclear envelope, (ii) an F-actin network supporting chromosome congression, and (iii) cortical contraction waves driving bulk cytoplasmic flows.

Of the three, the F-actin shell is the so far least characterized one. It has only been studied in detail in starfish oocytes in our earlier works^31,32^. Here, we show that in *Clytia* oocytes a very similar F-actin shell forms, and facilitates nuclear envelope disassembly as it does in starfish oocytes. In both species, the F-actin shell forms transiently on the inner side of the nuclear envelope at NEBD, persisting just for 1-2 minutes. The shell initiates from foci and then spreads in a wave-like manner engulfing the entire nucleus. By depleting available actin monomers, we were able to effectively block the formation of the F-actin shell. Then, by using dextran entry as a functional assay we show that the F-actin shell facilitates the rupture of nuclear membranes -- a nearly identical manner to starfish oocytes. In starfish, we were also able to visualize the ultrastructure of the F-actin shell showing that it extends F-actin spikes from the lamina into nuclear membranes. Whether these features are conserved to *Clytia* remains to be addressed in future work. Similarly, in starfish we showed that the F-actin shell is nucleated by the Arp2/3 complex, which needs to be confirmed in *Clytia*.

However, based on the striking similarity, we can already conclude with confidence that the transient F-actin shell facilitating nuclear envelope rupture is not an exotic feature of echinoderms, but a more broadly conserved mechanism of metazoan oocytes. Indeed, while characterized in much less detail, an F-actin shell along the nuclear envelope has been observed in other species, including the early embryos of the other cnidarian model *Nematostella vectensis*^55^, other echinoderm species, such as sea urchin^56^, in polychaete worms^57^, and somewhat similar structures have recently been reported in early mouse embryos, too^58^.

While speculative at this point, these observations together suggest that the F-actin shell is an adaptation evolved to rapidly clear the large and stable nuclear envelope of animal oocytes and early embryos, allowing cytoplasmic microtubules access to chromosomes. Extrapolating from findings in starfish oocytes, at the molecular level the F-actin shell may have evolved by relocation of the cortical Arp2/3 machinery -- responsible to generate filopodia and other cortical F-actin structures -- from the plasma membrane to the nuclear envelope. The identification of the molecular mechanisms, likely involving specific activators of Arp2/3 localized at the nuclear envelope, will be an exciting challenge for the future studies.

The second functional module is the F-actin network facilitating chromosome congression. Here, our work revealed an initial difference: in *Clytia* the nuclear F-actin network forms early during oogenesis and is already present in immature oocytes. By contrast, in starfish the F-actin network forms only upon NEBD. On the other hand, similar to *Clytia*, in immature *Xenopus* oocytes a nuclear F-actin network is present, caused by oocyte-specific downregulation of Exportin-6^17^. Our bioinformatic analyses identified that Exportin-6 expression is also downregulated in *Clytia* oocytes, suggesting a conserved mechanism. Thus, a nuclear F-actin network appears to be present in oocytes in some species, while it forms upon NEBD in others. Work in *Xenopus* oocytes suggest that the F-actin network functions to mechanically stabilize the nucleus^17^, and it also keeps nuclear bodies and nucleoli dispersed^46^. It will be interesting to see why in oocytes of some species such an F-actin network is functionally more important, while it is apparently not required in others.

In *Clytia*, the pre-existing, dense F-actin network disassembles after NEBD. Both speeding up disassembly (latrunculin B treatment) and slowing it down (phalloidin injection) interfered with chromosome congression and led to chromosome loss. Firstly, this demonstrates that the F-actin network is essential for chromosome congression and to prevent chromosome loss, as in starfish oocytes^13^ and in porcine and human oocytes^16^. Secondly, our findings in *Clytia* are consistent with a disassembly-driven contraction mechanism we first proposed in starfish^15^. Inhibitor treatments and computer simulations point to the importance of regulation of network disassembly rate; too fast or too slow disassembly both result in chromosome loss and ultimately the formation of aneuploid eggs. Addressing the underlying molecular mechanisms will be an exciting challenge for the future.

In starfish oocytes, ‘F-actin patches’ form around chromosomes that synchronize capture by microtubules^54^. These F-actin structures we did not observe in *Clytia*. Indeed, such F-actin patches may be less needed in *Clytia*, since the two microtubule asters move into the nuclear region, and are not anchored to the oocyte cortex like in starfish oocytes. Therefore, capture by microtubules can occur simultaneously without the need for synchronization. F-actin patches may thus be a less-well conserved adaptation possibly specific to starfish or echinoderm species.

Taken together, with our new findings in *Clytia* added, we conclude that nuclear F-actin networks commonly function in facilitating chromosome congression in oocytes across metazoan species^13,16,18^. As shown earlier, the oocyte nucleus, the germinal vesicle, is too large for microtubule asters to efficiently capture chromosomes scattered in this large nuclear volume^6,13^. Contraction of an extensive filament network, like a fishnet, is an adaptation perfectly suited for this purpose. Our data further indicate that the still poorly understood disassembly-mediated mechanism may be common to contraction of such fishnet-like networks collecting chromosomes in oocytes^15^.

Thirdly, we show that chromosomes first collected deep in the cytoplasm are transported by bulk cytoplasmic flows to the oocyte cortex. This process is fully dependent on Rho-mediated contractility as it is sensitive to C3-exoenzyme treatment, and is completely independent of microtubules. This process is reminiscent of mouse oocytes, in which chromosomes, together with the spindle, are also transported to the cortex^59^. While the flow patterns in mouse oocytes are strikingly similar to those we observed here in *Clytia*^59^, they have been proposed to be driven by a cortical Arp2/3 enrichment near the spindle^59^, whilst other studies suggest that the spindle is transported rather by myosin motors^21^. Neither of these proposed mechanisms has been validated by a physical model, and thus the exact mechanism remains unclear.

Our biophysical model provides a simple explanation for how such flow patterns emerge. Surface contraction waves are a common feature of oocytes first observed in *Xenopus*^34^. In recent years, the Bement and other laboratories established the detailed molecular mechanisms of how the broadly conserved Rho-pathway generates such waves in oocytes^60,61^, guided by cell cycle transitions^35,62^. Here, the only assumption we made is that the underlying cytoplasm is dragged along with the cortical contraction. As we show, implementing this into a biophysical model closely recapitulates the observed flow patterns, and simulated virtual tracers are transported similarly to chromosomes. Thereby, we demonstrate how the broadly conserved Rho-pathway -- that was already known to drive surface contraction waves in oocytes of diverse species -- can drive flows in the bulk cytoplasm, and that chromosomes can be transported by these flows. Notably, not all species take advantage of this mechanism: in starfish, surface contraction waves are present and they drive cytoplasmic flows^36^, but chromosomes are collected by cortex-anchored microtubules and thus flows are not involved in chromosome transport^54^. On the other hand, bulk cytoplasmic flows drive chromosome transport in *Clytia*, and the same mechanisms may be at work in mouse oocytes as well.

In summary, our data reveal multiple conserved mechanisms mediated by the actin cytoskeleton adapted to support the specialized meiotic divisions of oocytes. The existence of these mechanisms in an early-branching non-bilaterian species provides strong evidence that these mechanisms evolved early and are common to oocytes of animal species.

## Materials and Methods

### Clytia culture, maintenance and hatching

We obtained *Clytia hemisphaerica* female polyp colonies from the European Marine Biological Resource Centre’s (EMBRC) site at Villefranche-sur-Mer, France. We cultured and maintained polyp colonies in seawater aquariums at 18°C at the Animal Facility of the Max Planck Institute for Multidisciplinary Sciences (MPI-NAT), Göttingen Germany. *Clytia* jellyfish were hatched from polyp colonies in beakers exposed to light for 5 h. Freshly hatched *Clytia* jellyfish were kept in beakers with continuous stirring for 2 days and then transferred on day 3 first to a ‘nursery’ tank and then to kreisel tanks following established protocols^42^.

### Oocyte isolation, microinjection and maturation

4 weeks after hatching, fully grown *Clytia* jellyfish were maintained under a 16 h light / 8 h dark cycle. Adult *Clytia* were collected right before the dark cycle began and kept under light until further use. To isolate the oocytes, gonads were carefully dissected from the jellyfish using scissors and forceps the day before. Isolated gonads, suspended in filtered seawater (FSW), were then kept with the lights on overnight (16-20 h) in an incubator at 18°C to allow oocytes to fully develop. The following day, fully grown oocytes were isolated by carefully opening the gonad, and extracting them with help of a titanium wire loop, glass needle, and pipette suction. Isolated, fully grown oocytes were then placed in Petri dishes with FSW and kept up until the end of the day. For injection, oocytes were placed into Kiehart chambers prepared as described earlier for starfish oocytes^63^. Instead of mercury-filled needles, oocytes were microinjected using a Femtojet (Eppendorf) with constant-pressure. Oocytes were imaged typically 10-15 min after injection. Meiosis was induced by the maturation-inducing peptide WPRPamide^64^. NEBD typically occurred 10 min after WPRP addition.

### Recombinant protein expression and purification

Recombinant proteins were expressed in E. coli BL21(DE3) cells using a G031-based expression vector and selected on kanamycin-containing agar plates. Single colonies were used to inoculate 2YT precultures, which were grown overnight at 37°C and subsequently used to inoculate Terrific Broth (TB) medium supplemented with kanamycin and glycerol. Cultures were grown until an OD600 of approximately 0.8-1.0, after which the temperature was reduced to 25°C. Protein expression was induced at an OD600 of approximately 1.8-2.0 using 100-200 µM IPTG. Cultures were typically incubated overnight at 25°C following induction, whereas EB3-mVenus was induced for 4 h.

Bacterial cells were harvested by centrifugation and resuspended in a lysis buffer containing 20 mM Tris-HCl pH 8.0, 500 mM NaCl, 20 mM imidazole, 1 mM EDTA, and 20% glycerol. Cell suspensions were supplemented with 1-5 mM DTT and lysed by intermittent sonication. Clarified lysates were ultracentrifuged at 38,000 g for 1 h. 14xHis-tagged recombinant proteins were purified by Ni-NTA affinity chromatography. The resin was equilibrated in a lysis buffer and incubated with the clarified lysate. Following binding, the resin was washed first with buffer containing 20 mM Tris-HCl pH 8.0, 500 mM NaCl, 20 mM imidazole, 10% sucrose, 1 mM EDTA and 1 mM β-mercaptoethanol, followed by a second wash containing 20 mM Tris-HCl pH 8.0, 250 mM NaCl, 20 mM imidazole and 1 mM β-mercaptoethanol. Proteins were released by brSUMO-protease cleavage. Eluted fractions were analyzed by SDS-PAGE, and fractions containing the target protein were pooled and concentrated to ∼2 ml using 30- or 50-kDa molecular-weight-cutoff centrifugal concentrators. Concentrated samples were further purified by size-exclusion chromatography using an ÄKTA system equipped with a Sephadex 75 column. SDS-PAGE was used to analyze fractions corresponding to the major protein peaks, and the purest fractions were pooled. Where required, proteins were buffer-exchanged using PD-10 desalting columns into storage buffer containing 20 mM Tris-HCl pH 8.0, 250 mM NaCl, 1 mM β-mercaptoethanol, and 50 mM sucrose. Purified proteins were snap-frozen in liquid nitrogen and stored frozen at −70°C until use.

### Live cell fluorescent markers

Centrioles were visualized by expression of Poc1-ChemCherry (mCherry codon optimized to *Clytia*^47^). Poc1-ChemCherry was subcloned into a pGEM-HE-based vector containing the cyclin B1/B2 3′ UTR. The mRNA was synthesized *in vitro* from a linearized DNA template using the NEB HiScribe T7 ARCA mRNA Kit (with Tailing), followed by poly(A) tailing. The mRNA was purified using the Monarch RNA Cleanup Kit (NEB). Oocytes were microinjected with mRNA at 1.6 µg/µl concentration and incubated for 5 h to allow protein expression before imaging.

Chromosomes were labeled by microinjection of recombinant anti-histone nanobody^45^ tagged with mScarletI3 (4.2 mg/ml) or mVenus (7.15 mg/ml) protein. Microtubule plus ends were visualized by microinjection of purified EB3-mVenus (7 mg/ml) protein. F-actin was labeled using purified mScarletI3-UtrCH (6.93 mg/ml) or LifeAct-mScarletI3 (13.54 mg/ml) protein. To visualize the nuclear envelope and cellular membranes, DiIC18(5) oil (D307, Thermo Fisher Scientific) was prepared by diluting 25 mg DiIC_18_(5) oil in 1 ml of cooking oil and microinjecting it into oocytes. 500-kDa dextran conjugated to Abberior STAR RED was used as a cytoplasmic marker and co-injected with LifeAct-mScarletI3 in equal amounts. After protein injections, oocytes were incubated typically for 20 min at 18°C before imaging.

### Drug treatments

To fully stabilize F-actin, we first air-dried unlabeled phalloidin (20 µM) to remove the methanol solvent. Then resuspended the pellet in a buffer, injected it into oocytes, and incubated them for 20 min before imaging. To depolymerize microtubules, we diluted nocodazole to a 10 μM final concentration in FSW from a DMSO stock solution and added it to the chamber along with WPRP hormone. To depolymerize F-actin, latrunculin B was diluted to a 10 μM final concentration in FSW from a DMSO stock solution and added to the injection chamber along with the WPRP hormone. C3-exoenzyme powder was diluted in water (stock concentration, 104 µM), and 1 µL was loaded into an injection needle, injected into oocytes, and incubated for 20 minutes before meiosis induction. In the control groups, oocytes were administered either the same volume of DMSO solvent (at a final concentration of 1%) or were injected equal amounts of water, and incubated for the same durations of time.

### Immunostaining

Isolated oocytes were fixed at desired time points using a formaldehyde-based fixative (100 mM Hepes (pH 7.0, adjusted with NaOH or KOH), 50 mM EGTA, 10 mM MgSO4, 0.5% Triton X-100, 400 mM sucrose, and 1% formaldehyde)^65^. Oocytes were fixed for 1 h at room temperature, then washed in 1xPBS with 0.2% Triton X-100 under a microscope to avoid sample loss. After washing, samples were blocked in a blocking buffer containing 1xPBS, 0.1% Triton X-100, and 3% BSA. For antibody staining, the primary antibody was mixed with the corresponding secondary nanobodies or Fab fragments and incubated overnight at 4°C. Microtubules were labeled with an anti-α-tubulin mouse antibody (DM1A; Sigma-Aldrich), and nuclear pores with anti-FG-repeat antibody (mAb414; Sigma-Aldrich). Anti-mouse secondary nanobody donkey Fab anti-mouse IgG Alexa Fluor 594 (#715-587-003; Jackson) was used. F-actin was labeled with phalloidin conjugated to Alexa Fluor 647 (Abberior Star Red). Chromosomes were labeled with Hoechst 33342 (Thermo Fisher Scientific). Oocytes were mounted in PBS.

### Image acquisition and processing

Live and fixed oocytes were imaged using a Leica TCS SP5, Zeiss LSM880, or Zeiss LSM900 confocal microscope. For live imaging, the Leica TCS SP5 was used with an HC PL FLUOTAR 20x Dry NA0.5 or HCX PL APO CS 63x water NA1.2 objective, the LSM880 with a C-Apochromat W 40x NA1.2 objective, and the LSM900 with a LCI PlanNeofluar multi-immersion 63x NA1.3 objective. For 3D live imaging, time-lapse series of 10-30 optical sections (typically 1 µm apart, pinhole 1-1.5 Airy units) were acquired with time intervals of 5 s to 4 min. For 2D live imaging, single optical sections were acquired at 15-30 s intervals. Fixed oocytes were imaged using the LSM900 with the LCI PlanNeofluar 63x multi-immersion objective, or a Yokogawa W1 spinning-disk confocal system mounted on a Nikon Ti2 body and using PrimeBSI sCMOS cameras (Visitron Systems) with a CFI SR Plan Apo IR 60x NA 1.27 water immersion objective was used. For the LSM900, the pinhole was set to 1-1.5 Airy units, LSM900 images were deconvolved using the built-in ZEN LSMplus function. On the spinning disk system typical settings were: exposure time 100-500 ms, 10-25 z-slices with 3 or 4 μm step size.

### Chromosome tracking and spatial analysis

Anti-histone-nanobody-labeled chromosomes were tracked in confocal 3D time-lapse recordings acquired at 5 s intervals. Chromosomes and oocyte boundaries were segmented using a pixel-classification model in Ilastik (version 1.4.0)^66^. The model used background, foreground (signal of interest), and boundary classes, and the resulting probability maps were used to generate 3D label masks for the chromosomes and the oocyte boundary. Chromosome label masks were then inspected and manually corrected in Napari^67^ to retain individual or clustered chromosomes while excluding non-specific fluorescent objects. For each time point, an intensity-weighted centroid of the chromosome distribution was calculated within the chromosome label masks. The distance of each chromosome from this centroid was determined, and the mean distance to the intensity-weighted centroid was computed as a measure of chromosome spatial distribution. To quantify chromosome position relative to the oocyte cortex, the shortest distance between the chromosome centroid and the oocyte boundary was calculated at each time point and plotted over time. These analyses were performed for all experimental conditions using at least three independent datasets per condition.

### PIV analysis

To quantify cytoplasmic flows, the mitochondrial endogenous GFP signal was used, imaged in 2D along the midplane of the oocytes. Time-lapse recordings were first segmented, separating the oocyte from the outside background using simple thresholding, and this segmented label mask was then registered to correct for oocyte movements. This was done in FIJI/ImageJ^68^ using the Multistackreg and Turboreg plug-ins^69^. x and y coordinate shifts were then applied to the GFP and chromosome channels. After registration, recordings were analyzed by PIVlab (version 3.12)^70^ in MATLAB (version 25.2, R2025b, MathWorks). A mask was applied to exclude the nucleus and background from flow analysis. PIV analysis was performed using a fast fourier transform cross-correlation algorithm with two multipass windows of 64×64 and 32×32 pixels. We defined a velocity threshold for each flow field to exclude outliers. We extracted local PIV flow vectors based on chromosome positions. For this, chromosome tracking was done in Imaris (Oxford Instruments) using spot detection. Local flow near chromosomes was defined as a 30−30 μm square window. For speed correlation plots, a moving average (5 frames) of chromosome speeds was calculated; for local PIV vectors, no averaging was done.

### Bioinformatic analysis

*Clytia hemisphaerica* exportin-6 (EXP 6) sequences were retrieved from the MARIMBA database (Marine Invertebrate Models database; https://marimba.obs-vlfr.fr/), using a keyword search of the annotated protein dataset. All entries annotated as Exportin 6 were recovered and their homology was verified by BLASTP and TBLASTN searches, using NCBI non-redundant protein and nucleotide databases, respectively. The BLAST searches were run using the default parameters. The e-value threshold returned a value of 0.0 with a percentage identity of 100. Hits were considered orthologous on the basis of e-value, query coverage and best-hit recovery of XPO6 in the source organism. For comparative analysis, reference EXP 6 sequences were obtained from NCBI for *Clytia hemisphaerica* [XP_066934694.1] and *Xenopus laevis* [NP_001088605.1 xpo6.S; XP_041433775.1 xpo6.S]. Their identity was likewise confirmed by BLASTP and TBLASTN. Sequences were aligned using Clustal omega (CLUSTALW) online tool along with the default parameters, to assess conservation across orthologues. Domain architecture was annotated with InterProScan via the InterPro web server (https://www.ebi.ac.uk/interpro/ ; v106.0, accessed July 2025), submitting each protein sequence in FASTA format. The analysis was done with particular attention to functional domains and predicted nuclear export signals.

### Cytosim Simulations

Chromosome capture and positioning were simulated using Cytosim (version 3.10)^53^, a particle-based Brownian dynamics framework, in a 3D spherical domain of 60 µm diameter representing the nucleus. Microtubules were modeled as dynamic, semi-flexible polymers, nucleated from a mobile centrosome and exhibiting dynamic instability. Microtubule lengths were capped at 30 µm (maximum) and 0.05 µm (minimum). Centrosomes were introduced at the sphere surface with ∼23 µm spacing and initially confined to the surface for 1 s; thereafter, they were free to explore the interior volume. A soft spring-like boundary condition (spring constant:0.5 pN/µm) was applied only to segments extending beyond the sphere surface, preventing escape without imposing a hard wall.

10 chromosomes (4 µm diameter) were introduced randomly within a 6 µm shell adjacent to the boundary as observed in experiments. Each chromosome carried 10 actin-binding sites and 8 microtubule-binding sites. Chromosome-microtubule attachments were irreversible and represented stably bound dynein motors: processive, minus-end-directed motors, with a linear force-velocity relationship, v=v_0 (1-F/F_stall). The F-actin network was modeled under four distinct conditions to dissect the roles of contracting F-actin networks and microtubules on chromosome capture: Control: Actin filaments underwent disassembly at 0.0015 µm/s, with low medium viscosity (0.01 pN·s/µm²), enabling network contraction. Actin only: Actin filaments disassembled at 0.0015 µm/s, but microtubules were absent. Stable actin: Actin filaments were static (no disassembly), and medium viscosity was high (1 pN·s/µm²), mimicking a dense network and preventing network dynamics. Microtubules only: F-actin network was absent; microtubules were present. In all actin-containing conditions, filaments (rigidity: 0.075 pN·µm², initial length: 9 µm, discretized into 1.5 µm segments) were crosslinked by 120,000 end-tracking crosslinkers and 60,000 non-end-tracking crosslinkers. Both crosslinkers generated spring-like forces with a rest length of 0.01 µm and stiffness of 100 pN/µm. Additionally, microtubule plus-end crosslinkers were included to model experimental observations of correlated motion of the asters. The simulation ran for 400 s with a time step of 0.1 s. All parameters (Table S1) were chosen to reflect known biophysical properties and were consistent with prior modeling studies^15,53,54^.

### Cytoplasmic flow model

The model flow patterns were calculated from the changes in cortical F-actin fluorescence as follows: cortical F-actin dynamics were extracted from time-lapse fluorescence images. For each frame, an arbitrary reference point in the interior of the cell was chosen, and the cortex was divided into angular sectors. Within each sector, the radial position of the cortex outwards from the reference point was determined from the F-actin fluorescence. The contour was optimized to follow regions of high fluorescence while penalizing large changes in radius between neighboring sectors, thereby preventing the contour from following isolated intensity fluctuations. The penalty strength was tuned by visually inspecting the resulting contour. F-actin intensity was subsequently integrated in a narrow band around the obtained contour and divided by the sector-specific contour length to obtain the local cortical F-actin density.

To compare actin distributions between time points *t*, the contour was parametrized by normalized arclength *θ*, and profiles were mirror-symmetrized around the A-V axis of the cell. Assuming rotational symmetry around this axis, the measured F-actin density was converted into the zonal F-actin mass contained within surface zones of a sphere, accounting for the decreasing area of these zones toward the poles.

Movement of cortical F-actin between consecutive frames was estimated using optimal transport. In this approach, the actin mass is treated as a distribution that must be rearranged to produce the distribution in the next frame. Optimal transport determines the redistribution, the so-called transport plan, that achieves this while minimizing the total distance over which mass is moved. We calculated this transport using the one-dimensional Wasserstein distance implementation in the Python Optimal Transport library^71^. The resulting redistribution plan was used to calculate the mean displacement of actin at each surface position. Dividing this displacement by the time between frames provided an estimate of the local tangential cortical velocity *u_T_*.

The inferred cortical velocity was then used as the boundary condition for a spherical low-Reynolds-number flow model. We expanded the measured surface velocity into *N* standard axisymmetric squirmer modes

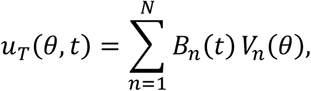

where *B_n_* describes the strength of each mode, and *V_n_* is defined from the corresponding Legendre polynomial *P_n_* as

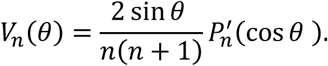

The coefficients *B_n_* were obtained by fitting this expansion to the cortical velocity inferred from optimal transport. The flow inside the cell was then calculated from these surface modes using the interior solution of the incompressible Stokes equations inside a sphere^72^. This is the internal analogue of the classical squirmer model and describes how tangential motion at the surface generates flow throughout a spherical volume. For each mode, the radial and tangential flow components were

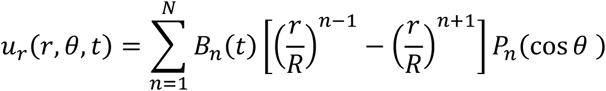

And

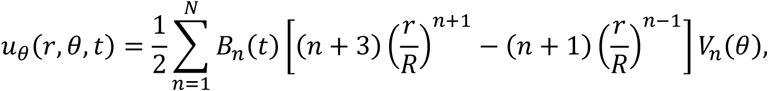

where *R* is the cell radius. These equations satisfy zero radial flow at the cell surface while reproducing the tangential cortical meridional velocity inferred from the change in actin fluorescence. Summing the fitted modes therefore yielded a three-dimensional, time-resolved estimate of the cytoplasmic velocity field inside the cell.

## Acknowledgements

We thank all members of the Lenart laboratory for protocols, reagents, and support, in particular Jasmin Jakobi for support with *Clytia* oocyte isolation and microinjection. We also thank the MPI-NAT Animal Facility, Sascha Krause, Sabrina Duthel, Tamina Hirschmeier, Daniela Wollrandt, Thomas Gundlach, Laura Kaulich, Philipp Missberger and Ulrike Teichmann, for their support. We thank the European Marine Biological Resource Center (EMBRC-France) for providing *Clytia* cultures, specifically the support of Laure Verdier and Axel Duchene. We thank Evelyn Houliston for her generous support and guidance with *Clytia* handling and protocols, and Tsuyoshi Momose for help with culture maintenance and for providing the Poc1-ChemCherry plasmid. We thank Trevor Huyton from Dirk Görlich’s department (MPI-NAT, Göttingen, Germany) for help with cloning, protein purification and for providing the Gibson assembly kit reagents. We thank Kevin Rentsch from Melina Schuh’s department (MPI-NAT, Göttingen, Germany) for providing us with pGEMHE-EB3(m)-mClover3 and pGEMHE-hMAD2L1-mVenus plasmids. We thank Emily Klass and Anne Wald (Institute for Numerical and Applied Mathematics, Applied Mathematics in the Natural Sciences, University of Göttingen) for discussions on flow field computation.

## Data and code availability

Image analysis scripts and biophysical models are available upon request.

## Author contributions

Yamini Vadapalli: conceptualization; formal analysis; investigation; visualization; methodology; writing (original draft); writing (review and editing). Komal Bhattacharyya: Cytosim model development and performing simulations. Sascha Lambert: flow model development. Bishal Samanta: bioinformatics analyses. Antonio Z. Politi: development 3D tracking workflow. Stefan Klumpp: resources and supervision of biophysical model development; funding acquisition. Peter Lenart: conceptualization; supervision; formal analysis; funding acquisition; writing (review and editing).

