## Supplementary material for "Cnidarian oocytes reveal conserved actin-driven mechanisms of female meiosis": Table S1

**Table S1: List of all parameters used in Cytosim simulations**

| **Component** | **Parameter** | **Value** | **Description** |
| --- | --- | --- | --- |
| **Geometry** | Sphere diameter | 60 µm | measured |
| **Microtubules** | Number per centrosome | 350 |  |
|  | Bending rigidity | 20 pN·µm² |  |
|  | Growth rate | 1.0 µm/s | Based on ref. 54, optimized |
|  | Shrinkage rate | –1.4 µm/s | Based on ref. 54, optimized |
|  | Catastrophe frequency | 0.2 s⁻¹ | Based on ref. 54, optimized |
|  | Max length | 30 µm | Length cap (l measured) |
|  | Min length | 0.05 µm |  |
|  | viscosity | 0.001 pN·s/µm² | Optimized |
| **Centrosome** | Radius | 0.02 µm |  |
|  | Viscosity | 0.001 pN·s/µm² | Optimized |
| **Microtubule plus-end crosslinkers** | Number | 500 |  |
|  | Binding rate | 10 s⁻¹ |  |
|  | Binding range | 0.25 µm |  |
|  | Unbinding rate | 10 s⁻¹ |  |
| **Chromosomes** | Number | 10 |  |
|  | Diameter | 4 µm |  |
| **Actin-binding sites** | Number per chromosomes | 10 |  |
|  | Binding rate | 50 s⁻¹ |  |
|  | Binding range | 0.15 µm | Ref. 15 |
|  | Unbinding rate | 0.0001 s⁻¹ | Ref. 15 |
| **Microtubule-binding sites** | Number per chromosome | 8 |  |
|  | Binding rate | 500 s⁻¹ |  |
|  | Binding range | 4 µm |  |
|  | Unbinding rate | 0 | Ref. 54 |
| **Dynein** | Unloaded speed (v_0_) | 1 µm/s |  |
|  | Stall force (F_stall_) | 100 pN |  |
|  | Force-velocity relation | v = v_0_(1 - F/F_stall_) | Linear relationship |
| **Actin filaments** | Rigidity | 0.075 pN·µm² |  |
|  | Initial length | 9 µm |  |
|  | Segment length | 1.5 µm |  |
|  | Number | 2,000 |  |
|  | Disassembly rate | 0.0015 µm/s | Ref. 54 |
| **Crosslinkers** | End-tracking | 120,000 |  |
|  | Binding rate | 50 s⁻¹ | Optimized |
|  | Binding range | 0.08 µm | Optimized |
|  | Unbinding rate | 0.01 s⁻¹ | Ref. 54 |
|  | Non-end-tracking | 60,000 |  |
|  | Binding rate | 50 s⁻¹ | Optimized |
|  | Binding range | 0.05 µm | Optimized |
|  | Unbinding rate | 0.01 s⁻¹ | Ref. 54 |
|  | Spring rest length | 0.01 µm | Ref. 54 |
|  | Spring stiffness | 100 pN/µm | Based on ref. 54, optimized |
| **Medium** | Viscosity | 0.01 pN·s/µm² | Optimized for contraction |
|  | Viscosity (stable actin) | 1 pN·s/µm² | Optimized for stable network, mimicking high filament density |
| **Simulation** | Time step | 0.1 s | Optimized |
|  | Duration | 400 s |  |
